# Input-dependent modulation of MEG gamma oscillations reflects gain control in the visual cortex

**DOI:** 10.1101/188326

**Authors:** Elena V. Orekhova, Olga V. Sysoeva, Justin F. Schneiderman, Sebastian Lundström, Ilia A. Galuta, Dzerasa E. Goiaeva, Andrey O. Prokofyev, Bushra Riaz, Courtney Keeler, Nouchine Hadjikhani, Christopher Gillberg, Tatiana A. Stroganova

## Abstract

Gamma-band oscillations arise from the interplay between neural excitation (E) and inhibition (I) and may provide a non-invasive window into the state of cortical circuitry. A bell-shaped modulation of gamma response power by increasing the intensity of sensory input was observed in animals and is thought to reflect neural gain control. Here we sought to find a similar input-output relationship in humans with MEG via modulating the intensity of a visual stimulation by changing the velocity/temporal-frequency of visual motion.

In the first experiment, adult participants observed static and moving gratings. The frequency of the MEG gamma response monotonically increased with motion velocity whereas power followed a bell-shape. In the second experiment, on a large group of children and adults, we found that despite drastic developmental changes in frequency and power of gamma oscillations, the relative suppression at high motion velocities was scaled to the same range of values across the life-span.

In light of animal and modeling studies, the modulation of gamma power and frequency at high stimulation intensities characterizes the capacity of inhibitory neurons to counterbalance increasing excitation in visual networks. Gamma suppression may thus provide a non-invasive measure of inhibitory-based gain control in the healthy and diseased brain.

## Introduction

Cortical gamma-band oscillations arise from the interplay between neural excitation (E) and inhibition (I)^1^ and may provide a non-invasive window into the state of the E-I balance in human cortical networks^2–5^. Being initially described in the visual cortex of cats in response to motion of high-contrast gratings^6^, gamma oscillations can be induced by similar visual stimuli in humans and measured with magneto-encephalography (MEG)^7^. While recent studies have questioned the potential MEG gamma parameters have for reflecting the inner state of E-I circuitry when they are measured in a single experimental condition^8^, the *modulation* of gamma oscillations by intensity of sensory inputs may help to characterize networks’ excitatory state through probing its input–output gain.

In the sensory cortex, the power of gamma oscillations is low in the absence of sensory stimuli or when the stimulation intensity is low^9–11^. The frequency of gamma oscillations increases monotonically with increasing intensity of visual input, such as visual contrast^9,12–14^ or motion velocity/temporal frequency^15,16^. Experimental studies in animals suggest that an increase of the tonic excitability of inhibitory interneurons may account for the stimulation-driven increase in gamma frequency^17,18^. On the other hand, the power of gamma oscillations in local field potentials (LFP) of non-human primates initially increases with contrast ^9,13,14^ or temporal frequency^19^ and then attenuates at higher stimulation intensities. This bell-shaped input-output relationship can be explained by an initially facilitative influence of rising excitatory drive on gamma synchrony in E-I circuitry^9,20^, followed by a suppressive effect (at high input intensity) that is mediated by an excessive recurrent inhibition from I-cells that desynchronizes E-cells’ firing^21,22^. If replicated in humans, such non-linear gamma dynamics may be a valuable measure that reflects the capacity of inhibitory circuitry to regulate the network's excitatory state.

Previous MEG studies in humans include modulation of the intensity of the visual input by varying either contrast or motion velocity. In the case of increasing visual contrast of static gratings, a near-linear facilitative effect on MEG gamma synchronization persists up to maximal contrast^12,23^. It is likely, however, that the bell-shaped changes in human MEG gamma power are only evident when the visual input is driven with higher stimulation intensities. While manipulations with static stimuli are limited in their capacity to increase the input drive, this can be achieved by the varying velocity/temporal frequency of moving high-contrast gratings. Indeed, because studies that used different stimulation parameters reported either velocity-related increases^24^ or decreases^25^ in gamma power, the effect of velocity on MEG gamma response might be nonlinear and resemble that which has been observed in the LFP of primates^19^.

We predicted that modulating excitatory drive to the cortex (by increasing the velocity/temporal frequency of visual motion) would result in a monotonic increase of MEG gamma peak frequency and concurrent bell-shaped change in gamma power. We further hypothesized that the reduction of MEG gamma response at high stimulation velocities would reflect the capacity of highly excited inhibitory neurons to limit increasing excitation in the whole visual circuitry. As increasing excitation of inhibitory neurons leads to acceleration of gamma oscillations^17,18^ this explanation predicts that greater increases in gamma frequency would be associated with a stronger suppression of gamma amplitude at the descending branch of the bell-shaped curve.

Developmental changes of excitatory and inhibitory neurotransmission^26–28^ leave a possibility that our previous observation of motion-velocity modulation of MEG gamma oscillations in children^25^ is age-specific. To clarify this issue, we investigated a large group of subjects spanning ages from 7 to 40 years. In the first experiment, we used static and moving stimuli to study the theoretically-predicted dependencies of gamma power and frequency on the velocity of visual motion. In the second experiment, we focused on the suppressive effect of high motion velocities on induced gamma oscillations, and tested for the hypothesized relationship between velocity-related modulations of gamma frequency and power.

## Results

### Experiment 1

In the 1^st^ experiment, participants observed annular gratings that were moving at velocities of 0 (static), 1.2 (slow), 3.6 (medium), and 6.0 (fast) °/s. Each trial included a stimulus presentation for 1200–3000 ms, followed by a blank screen with a fixation cross (see Methods for details). All participants showed highly reliable sustained gamma response to the stimuli, with a monotonic increase of gamma oscillation frequency as a function of visual motion velocity. The group mean gamma frequencies were 51.1, 56.5, 65.3, and 70.5 Hz for the static, slow, medium and fast conditions, respectively (Fig. 1C). No correlation was found between the peak gamma frequency measured at each of the 4 velocity conditions and velocity-related changes in frequency (16 correlations, all p's>0.1). This contradicts the idea that the peak frequency of gamma response ‘saturates’^29^ at the motion velocities used in this and previous studies.

Unlike the changes in frequency related to visual motion velocity, the changes in gamma power were clearly non-linear. Nearly all (9 of 10) subjects demonstrated an initial increase in gamma power for the slow velocity, followed by power suppression at the medium velocity. The further increase in motion velocity (to fast) resulted in gamma suppression in all 10 participants (Fig. 1E). Thus, the bell-shaped dependency of gamma amplitude as a function of motion velocity was observed in nearly all subjects.

**Figure 1.**
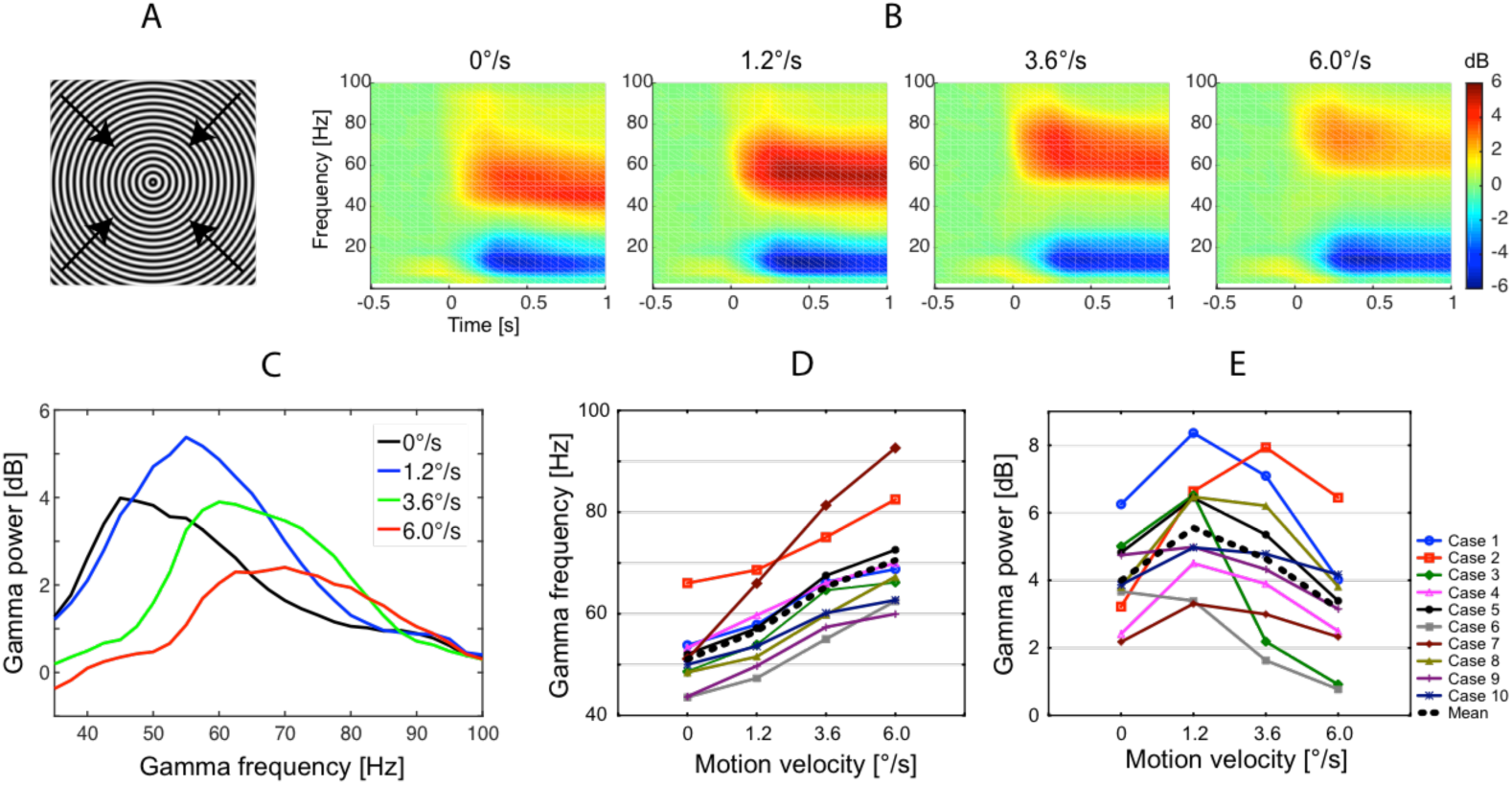
Changes in gamma power and frequency induced by the visual presentation of the annular gratings. The grating displayed in panel A either remained static (0°/s) or drifted at one of the three velocities (1.2°/s, 3.6°/s or 6.0°/s) in the direction indicated by the arrows. B. Grand average time-frequency plots. C. Grand average spectra of sustained gamma response. Gamma power was computed in the 200 - 1200 ms time interval after stimulus onset and was normalized to the pre-stimulus baseline (−900 to 0 ms). D-E: Individual peak frequency (D) and power (E) of sustained gamma responses. Here and hereafter, the frequency range of interest was established where the post-/pre-stimulus power ratio exceeded 2/3 of the peak value. The center of gravity of the power over this frequency range was used as the gamma peak frequency. The average power over these frequencies was used as the gamma peak power (see Methods for details).

The suppression of gamma power with increasing temporal frequency is unlikely to be driven by a dampened neural response to a non-preferable temporal frequency. Indeed, according to studies in non-human primates, most neurons in V1 respond well at temporal frequencies up to 6–10.0 Hz (medium to fast velocity in our study) and their responses then drop at higher frequencies^30–32^. Unlike the neural firing rate, V1 gamma oscillations in monkeys begin to attenuate already after 2 Hz^19^, which corresponds to the slow velocity in our study. The fMRI study of Singh et al^33^ also undermines the role of dampened neural activation in the gamma response suppression observed in the current study. These authors used high-contrast gratings with spatial frequency of 2 cycles per degree of visual angle (i.e., similar to ours that had 1.66 cycles per degree). Their gratings drifted with temporal frequencies of 0, 2, 5, 9, and 18 Hz and they observe *maximal fMRI activation* in visual cortical areas for the 9 Hz stimulus, i.e. the one with a temporal frequency closest to that of our ‘fast’ stimulus. Since fMRI BOLD signal correlates *positively* with neuronal spiking^34^, the results of Singh et al suggest that the gamma response suppression occurs at the peak of neuronal activation.

An alternative explanation for the suppression of gamma power is based on the direct relationship between gamma frequency and excitability of inhibitory neurons^17,18^. A monotonic increase in gamma frequency with increasing motion velocity in both humans (current study) and animals^15,35^ points to a growing tonic excitability of inhibitory neurons. This, in turn, may be a crucial factor that mitigates excessive excitation while suppressing gamma oscillations^21,36,37^.

A computational study by Borgers and Kopell (2005) suggests that the suppressive effect of over-excited inhibitory cells on gamma oscillation is lower when the E/I ratio is higher^21^. The magnitude of gamma suppression at high motion velocities may thus help to characterize the E-I balance in the visual cortex. We therefore investigated this suppressive effect in more detail with a second experiment that focused on inter-individual variability, the predicted correlation with frequency changes, and developmental effects.

### Experiment 2

In Experiment 2, participants were presented only with moving gratings, i.e. visual motion velocities of 1.2 (slow), 3.6 (medium) and 6.0 °/s (fast); the experimental setup was otherwise similar to that of Experiment 1. Figure 2 shows grand average time-frequency (A,B) and spectral (D,F) plots based on the results from 46 children and 26 adults.

**Figure 2.**
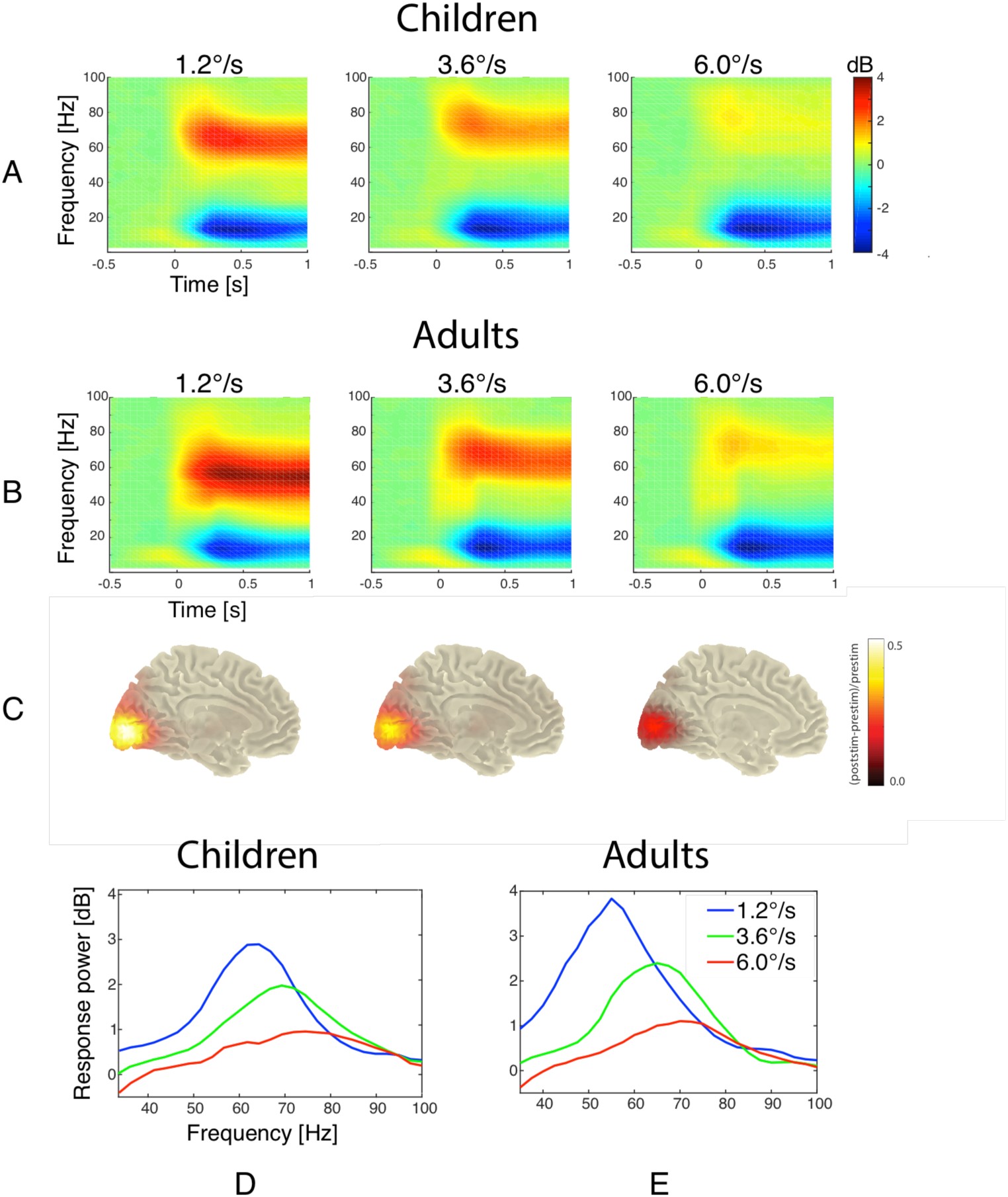
Gamma signals induced by visual presentation of annular gratings drifting at slow, medium, and fast velocities in children and adults. The two upper rows present grand averaged time-frequency plots for the ‘maximal pair’ of gradiometers in children (A) and adults (B). Source localization with DICS beamformer (C) is based on the adults’ data. A transparency factor was applied to mask values that are close to zero. The bottom row is grand averaged spectra of gamma responses for children (D) are adults (E).

#### Incidence of the reliable gamma responses

Table 1 shows the number of subjects who demonstrated stimulus-related increases in gamma power at p<0.05 and p<0.0001 probability levels. Under the slow and medium velocity conditions, gamma response peaks were highly reliable (response vs. baseline: p<0.0001, see Methods for details) in all adults and in the majority of children. For the fast velocity, highly reliable peaks were detected in the majority of adults and in about half of the children.

**Table 1.** Incidence, mean frequency and mean power of motion-related gamma responses detected at different p-levels (Mann-Whitney U test) in children and adults in sensor space. Standard deviations are given in parentheses.

| P<0.05 |  |  |  | P<0.0001 |  |  |
| --- | --- | --- | --- | --- | --- | --- |
| Children, N=46 |  |  |  |  |  |  |
| Stimulus velocity | Frequency, Hz | Power, dB | N (%sample) | Frequency, Hz | Power, dB, | N (%sample) |
| Slow | 66.1(6.1) | 3.1(2.0) | 45 (98%) | 65.2 (5.3) | 3.6 (1.8) | 37 (80%) |
| Medium | 73.2(8.5) | 2.2(1.8) | 43 (94%) | 72.3 (8.0) | 2.8 (1.8) | 31 (67%) |
| Fast | 82.2(10.8) | 1.3(1.0) | 37 (80%) | 79.6 (7.4) | 1.9 (1.1) | 21 (46%) |
| Adults, N=26 |  |  |  |  |  |  |
| Stimulus velocity | Frequency, Hz | Power, dB | N and %sample | Frequency, Hz | Power, dB | N and %sample |
| Slow | 55.7(5.7) | 3.8(1.4) | 26 (100%) | 55.7(5.7) | 3.8(1.4) | 26 (100%) |
| Medium | 65.1(6.0) | 2.6(1.3) | 26 (100%) | 65.1(6.0) | 2.6(1.3) | 26 (100%) |
| Fast | 70.0 (8.5) | 1.3 (0.7) | 26 (100%) | 69.5 (5.4) | 1.5 (0.7) | 19 (73%) |

#### Sensor- vs. source-derived gamma responses

The high Spearman correlations between source- and sensor-derived gamma response power (0.79 to 0.91, all p’s<0.0001) and frequency (0.83–0.96, all p’s<0.0001) suggest a close correspondence between sensor- and source-derived results. These findings agree well with those of Tan et al^38^ who reported high reliability of both source- and sensor-derived parameters of the visual gamma response and their close correspondence.

DICS (dynamic imaging of coherent sources) beamformer analysis^39^ showed that the mean position of the gamma response’s maximum in all three conditions corresponded to the same MNI template voxel in the left calcarine sulcus. We did not find evidence for a systematic shift of the response’ maximum position with changes in motion velocity. Our findings suggest that the same neural circuits participate in the generation of gamma activity in all three velocity conditions. Details on source analysis are presented in *Supplementary information*.

Given the fact that structural MRI data were not available for all of the participants, we restricted further analyses to the sensor data.

#### Velocity-related modulations of gamma response, their variability and outliers

<u>Power.</u> To analyze the effect of visual motion velocity on the gamma response power at the group level, we included only those participants who demonstrated reliable (p<0.0001, see Methods for details) gamma peaks under the slow condition. This strict criterion did not affect the size of the adult sample, but reduced the number of children in the analysis (adults: 26 of 26, children: 37 of 46).

**Figure 3.**
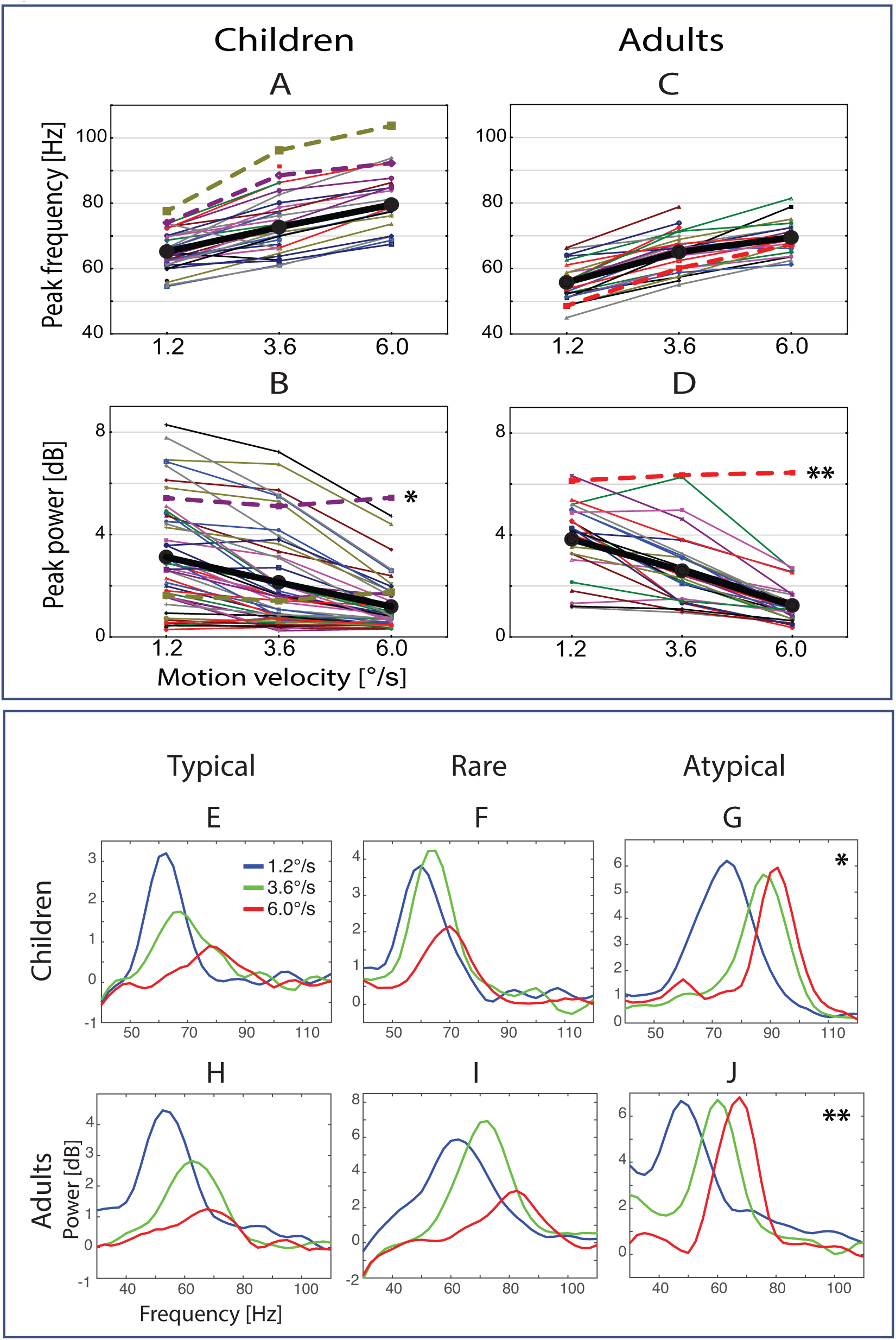
Individual variability of gamma response parameters. *A,B,C,D:* Individual gamma peak frequency and power values under the three velocity conditions in children and adults. Thick black lines show group mean values. Dashed lines refer to the subjects (2 children, 1 adult) with an atypical lack of gamma suppression at the highest velocity. *E, E, G, H, I, J:* Power spectra of representative subjects. E, H: Typical examples of gamma suppression with increasing motion velocity observed in children and adults. F,I: Rare variants of gamma response with an initial increase in gamma power at the medium velocity followed by power suppression at the fast one. G,J: An atypical absence of gamma suppression at the fast stimulus velocity. The subjects with atypical gamma responses displayed in G and J are marked by asterisks in panels B and D. See text for details.

The majority of participants, regardless of their age, demonstrated gradual attenuation of gamma response with increasing stimulus velocity from slow, through medium, to fast velocity conditions (Fig. 3E,H). A few subjects - three adults (11% of adults) and two children (5% of children) - showed an initial increase in gamma power at the medium velocity condition followed by its suppression at high velocity (Fig. 3F,I; see also Fig. 1E). One adult (Fig. 3J) and two children (one of them is presented in Fig.3G) displayed virtually no velocity-related changes in gamma response power, even at the high velocity. Visual inspection of the MEG recording of the adult subject revealed spike-and-wave complexes over the posterior brain regions. Post-hoc examination indicated that he had an unreported epilepsy diagnosis. Two children who displayed an atypical lack of gamma suppression also had medical conditions that were revealed post-hoc (see Methods for more information about these participants). These three cases were excluded from all further group analyses.

To analyze the effect of velocity on gamma response power we included only those subjects who had reliable gamma responses in at least one of the conditions (37 children and all 26 adults). Repeated measures ANOVA results supported a profound reduction of gamma response power with transition from the slow to the medium and further to the fast condition in children (F_(2,72)_=93.0, ε=0.8, p<10e^−6^; η^2^=0.73) and adults (F_(2,50)_=83.1, ε=0.83, p<10e^−6^; η^2^=0.77).

To analyze the effect of velocity on gamma frequency we included those subjects who had reliable gamma responses in all three experimental conditions (21 children and 19 adults). There was a robust effect of visual motion velocity on gamma frequency in both children (F_(2,40)_=173.6, ε=0.8, p<10e^−6^; η^2^=0.9) and adults (F_(2,36)_=145.9, ε=0.8, p<10e^−6^; η^2^=0.89). The average increase in gamma frequency from the slow to the fast condition was 15.3 Hz in children and 14.6 Hz in adults.

#### Effect of age on gamma oscillations

*<u>Power.</u>* The power of the stimulus-induced gamma response significantly increased with age in children between 6 and 15 years, but did not change in adults between 19 and 40 years (Table 2; Fig. 4A).

**Figure 4.**
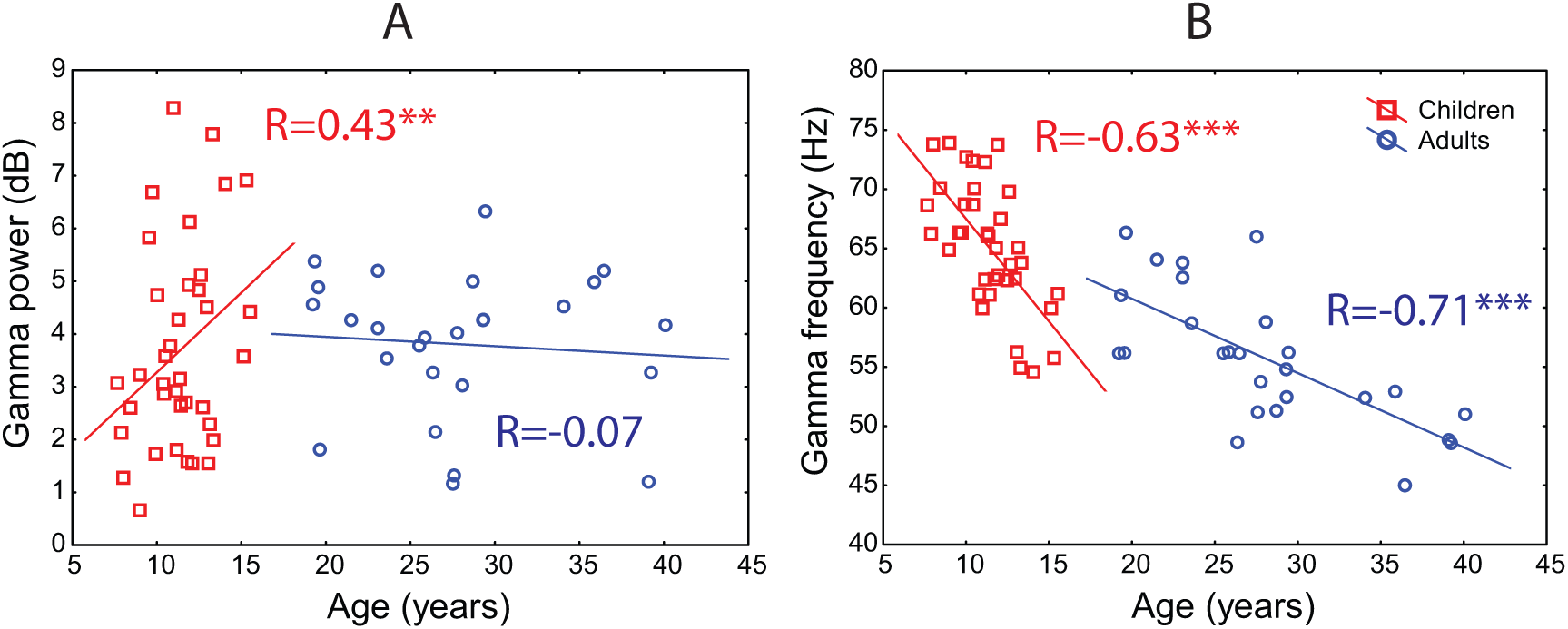
Developmental changes of gamma response power (A) and frequency (B) in children and adults under the ‘slow’ velocity condition. (Spearman correlations, **p<0.01; *** p<0.0001).

**Table 2.** Correlations of gamma parameters with age in children and adults (Spearman’s R).

|  |  |  |
| --- | --- | --- |
| Stimulus velocity | Children | Adults |
| Amplitude |  |  |
| Slow | **0.43\*\***(N=46) | -0.07(N=26) |
| Medium | **0.43\*\***(N=46) | -0.13(N=26) |
| Fast | **0.37\***(N=46) | -0.30(N=26) |
| Frequency<sup>#</sup> |  |  |
| Slow | **-0.63\*\*\***(N=37) | **-0.71\*\*\***(N=26) |
| Medium | **-0.39\***(N=31) | **-0.74\*\*\***(N=26) |
| Fast | **-0.54\***(N=21) | -0.42(N=19) |

<sup>#</sup>Frequency was assessed in case of reliable gamma response peak ( $p < 0.0001$ , see Methods for details). Spearman correlations, \* $p < 0.05$ ; \*\* $p < 0.01$ ; \*\*\* $p < 0.0001$

ANOVA with the main factor Age-Group (adults, children) and repeated measures factor Velocity (3 conditions) revealed a highly significant effect of Velocity on gamma power (F_(2, 122)_=172.8, ε=0.87, p<1e^−6^; η^2^=0.74), but no effect for the Age-Group or the Age-Group*Velocity interaction (p’s>0.6). This means that the gamma suppression magnitude remained essentially the same across childhood and adulthood.

*<u>Frequency.</u>* Gamma peak frequency was assessed only when the gamma peak was determined to be reliable (p<0.0001, see Methods for details). In both children and adults, the gamma peak frequency decreased with age (Table 2; Fig. 4B). Notably, the drop in gamma frequency was more rapid during childhood (1.71 Hz per year for the slow condition) than during adulthood (0.64 Hz per year for the slow condition). To test for a difference in the rates of gamma frequency changes in children and adults, we compared the respective regression coefficients. A homogeneity of slopes analysis confirmed a significant difference between gamma frequency age-related slopes in children and adults for the ‘slow’ (F_(1, 59)_=7.8, p<0.005, η^2^=0.13) and the ‘fast’ (F_(1, 36)_=7.5, p<0.01, η^2^=0.17) velocity conditions.

ANOVA with factors Age-Group (adults, children) and Velocity (3 conditions) confirmed highly significant effects of both Age-Group (F_(1, 38)_=27.1, p<0.0001; η^2^=0.42) and Velocity (F_(2, 76)_=317.2, ε=0.81, p<1e^−6^; η^2^=0.89) on gamma frequency. Inspection of the respective means showed that, on average, gamma frequency was higher in children aged 7–15 years than in adults (64.3, 72.4, and 79.9 Hz in children vs 54.8, 63.9, and 69.5 Hz in adults for the slow, medium and fast velocities, respectively). However, the lack of an interaction between Age-Group and Velocity (p=0.4) suggests that the magnitude of velocity-related increase in gamma frequency is developmentally stable.

#### Effect of increasing excitatory drive on gamma suppression

Speeding up the motion of the drifting gratings increases the intensity of the visual input and is likely to result in a stronger excitatory drive to the visual cortex. To estimate changes in the excitatory drive, we measured pupil reactions in a subset (19 of 26) of our adult participants. Pupil constriction can be induced by changing the attributes of a visual stimulation (e.g. spatial structure, color, or motion coherence) without any changes in light flux^40–42^. This phenomenon is thought to reflect the activation of brain structures involved in the processing of visual features^40^. We therefore hypothesized that the presence of a velocity-related increase in pupil constriction would reflect rising activation of the visual cortical areas, including V1.

The repeated measures ANOVA revealed a highly reliable effect of Velocity (slow, medium, and fast) on pupil response (F_(2, 36)_=17.3, e =0.92, p<0.0001; η^2^=0.49; Fig. 5). The magnitude of pupil constriction increased with increasing motion velocity of the drifting grating. Apart from a strong pupillary constriction effect, the fast motion was rated by the participants as the most intense and unpleasant condition; this also pointed to the strong excitatory drive produced by the fast stimulation.

**Figure 5.**
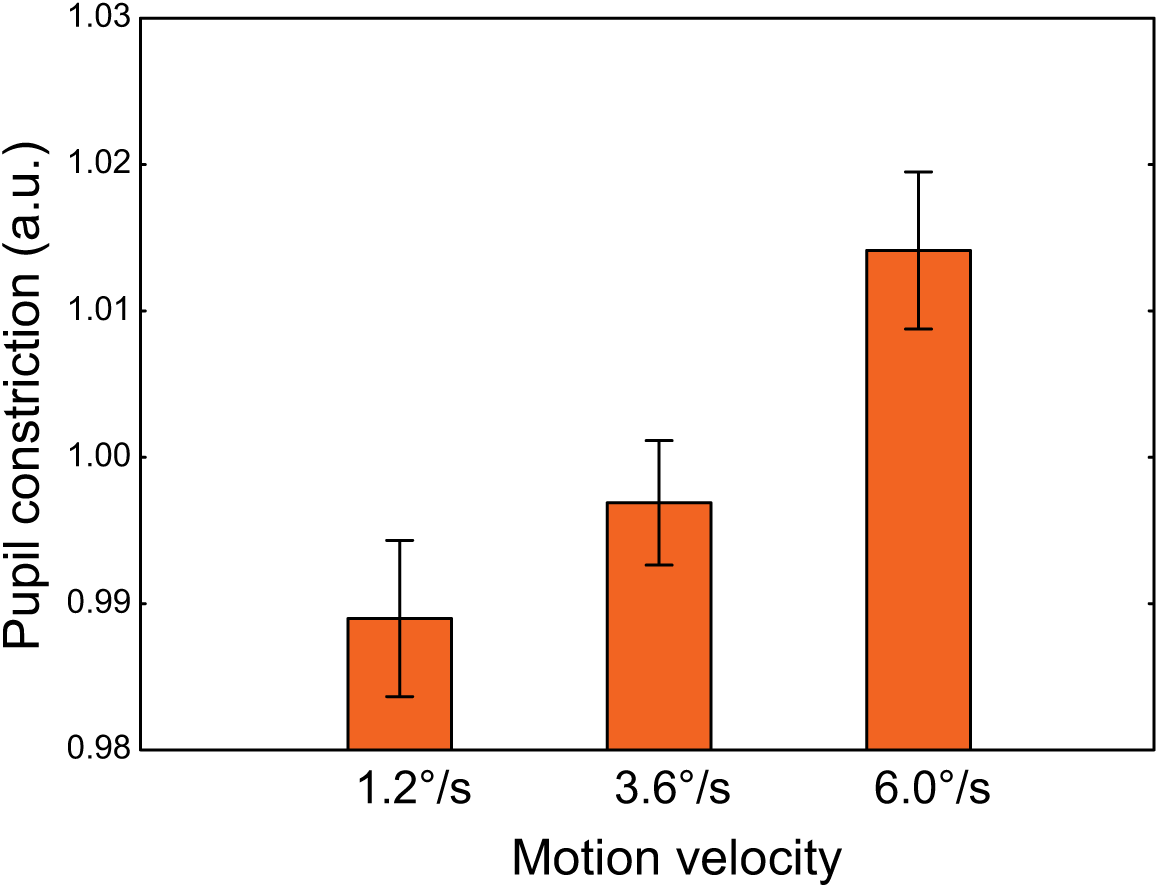
Pupillary constriction response to the annular gratings moving at the three velocities. Vertical bars denote 95% confidence intervals. The pupil constriction values were z-transformed across conditions for each subject.

The strong excitatory drive may cause both an increase in gamma frequency^17,18^ and suppression of gamma power^36^ through increasing excitation of inhibitory cells. Therefore, we predicted that a greater increase in gamma frequency with increasing velocity of visual motion would be accompanied by stronger reduction of the gamma power. To test this prediction, we analyzed the correlations between changes in gamma frequency and the corresponding normalized changes in gamma power between slow vs. medium and medium vs. fast velocity conditions. All participants with reliable (p<0.0001) gamma peaks under the corresponding conditions were included in this analysis. A greater increase in gamma frequency indeed correlated with a stronger suppression of gamma power from the slow to the medium condition (Spearman R_(66)_=0.32, p<0.01, Fig. 6). For the medium vs. fast condition, the correlation did not reach significance (R _(50)_=0.16, p=0.28).

**Figure 6.**
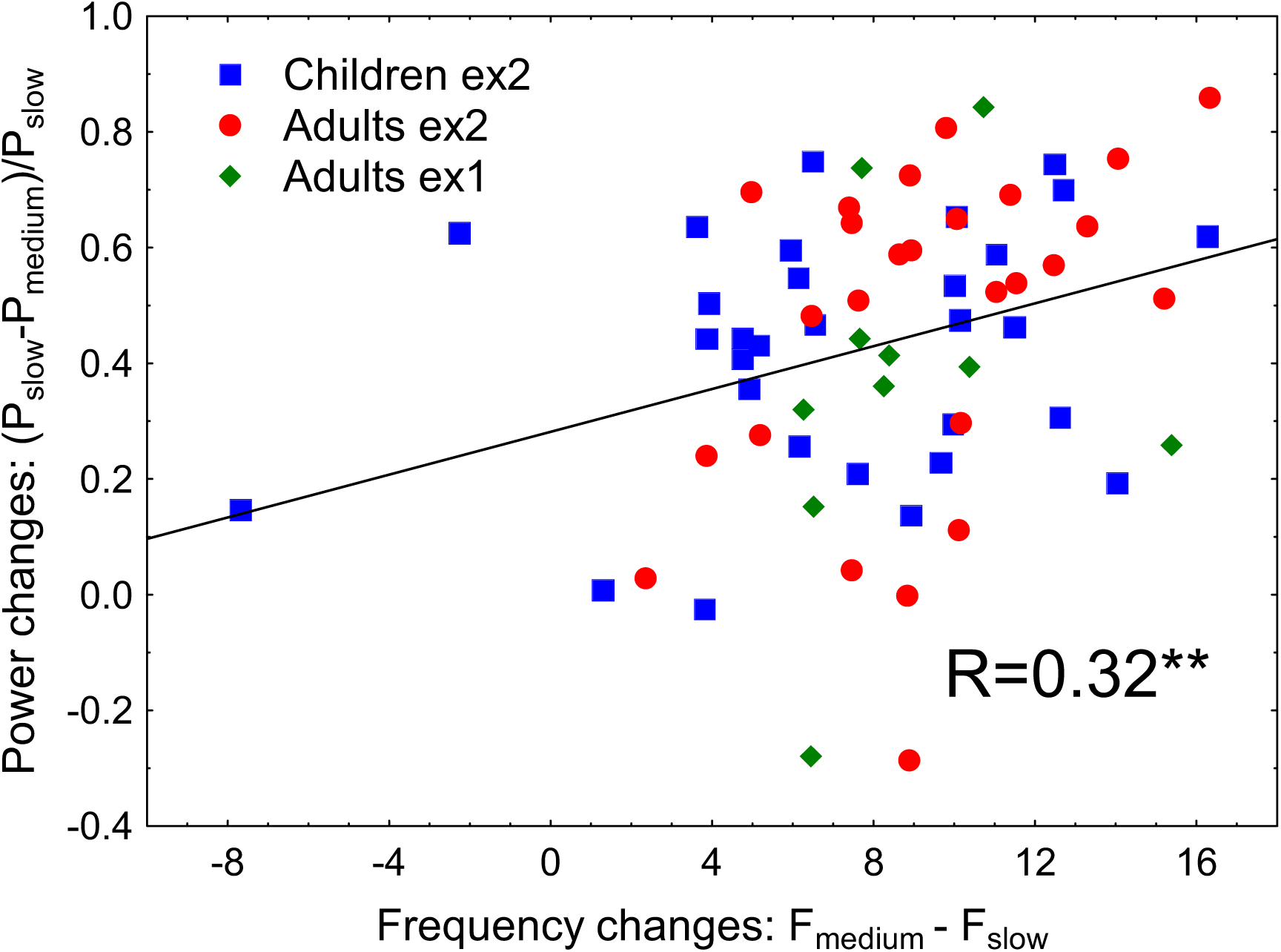
The relationship between changes in gamma frequency and normalized changes in gamma power induced by transition from the slow to the medium stimulus velocity. The regression line corresponds to the whole sample analysis; the color and shape of the individual values represent the respective experimental groups. Spearman correlation for the whole sample: **p<0.01.

## Discussion

We found that increasing the velocity of visual motion from 0 to 6°/s resulted in an approximately linear increase in frequency and clearly nonlinear changes in power of sustained gamma oscillations in MEG. The observed modulations were characterized by high magnitude and remarkable reliability. The same monotonic increase in gamma frequency with increasing velocity of visual motion has previously been observed in intracranial EEG of cats^15,35^ and in MEG recordings of children^25,43^. Here, we reproduced the MEG findings for a greater sample of participants (i.e. that comprised adults) and for a broader range of velocities (by including a static stimulus). The bell-shaped velocity-related changes in gamma power observed in the present study explain the seemingly contradictory findings that previously showed either velocity-related enhancement^24,29^ or suppression^25,43^ of gamma power. We also found that, despite drastic developmental changes in the frequency and power of gamma oscillations, the magnitude of their modulations by stimulus velocity remained relatively constant within a broad age-range of 7–40 years. Below, we discuss the neural basis for such velocity-related modulation of gamma oscillations, its relation to the E-I balance, as well as developmental changes in gamma response.

The modulations of MEG gamma oscillations by increasing velocity of visual motion observed in the present study (Fig. 1) mirror those found in LFP of animals in response to changes of temporal frequency/velocity^15,16,35^ and contrast^9,12,14^. In both humans and animals, gamma oscillations monotonically accelerate with increasing input intensity. Neurophysiological studies in animals suggest that the frequency of gamma oscillations is controlled mainly through changes in the tonic excitatory state of fast-spiking inhibitory cells^17,18^. We therefore suggest the effect of rising excitatory drive on MEG gamma frequency in the current study is mediated primarily through the increasing excitation of inhibitory neurons.

The initial increase in gamma power during the transition from the static to slowly moving gratings (Fig.1A,C,E) is likely caused by a transition from sub to near ‘optimal’ input intensity that resulted in recruitment of a large fraction of E and I cells in synchronous activity^9,20^. However, yet stronger excitation within both excitatory and inhibitory circuitry, triggered by a further increase in motion velocity, has a suppressive effect on gamma synchrony. Modeling studies suggest that excitation of inhibitory neurons beyond some ‘transition point’ leads to desynchronization in the cumulative activity of excitatory cells^21,36^ and thus results in suppression of gamma oscillations. The presence of this transition point could explain the bell-shaped input-utput relationship in the power of MEG gamma oscillations that we observed (Fig. 1C,E). The direct correlation between the rise in gamma frequency and the fall in power at the descending branch of the bell-shaped curve (Fig.6) fits well with this explanation: both gamma (power) suppression and gamma acceleration (frequency increase) are putatively caused by ‘over-excitation’ of inhibitory circuitry.

Notably, when the input intensity was modulated by increasing the *contrast* of static visual stimuli, previous studies demonstrated a gradual increase in gamma synchronization (i.e., power), without any evidence of gamma suppression at the highest contrast^12,23^. We suggest that the stimulation intensity attained with full-contrast stationary gratings is below the point of transition to MEG gamma suppression.

The Borger and Kopell’s model predicts that weaker gamma suppression at high intensities of excitatory drive occurs when inhibitory synaptic currents are less efficient at counterbalancing the growing excitation in the excitatory circuitry, i.e. in the case of a higher E/I ratio^21^. It is interesting in this respect that one of the three participants in our study who failed to suppress gamma in the fast velocity condition (Fig. 3J) had a post-hoc revealed diagnosis of epilepsy. The above considerations give hope to the idea that the modulation of gamma oscillations by velocity/temporal frequency of moving gratings may help to characterize the E-I balance in normal and abnormal visual networks. In the future, systematic studies specifically examining this issue in typical and atypical individuals may then have important clinical implications.

In the present study, we observed profound developmental changes in gamma response (Fig. 4; Table 2). These changes may reflect developmental alterations in GABAergic and glutamatergic neurotransmission that occur during childhood, adolescence and even adulthood^26–28,44–47^.

The developmental increase in the strength of the visual gamma response (Table 2, Fig. 4A) is compatible with previous reports of a maturational rise in auditory gamma oscillations evidenced by an increase in amplitude of 40-Hz auditory steady-state responses^48,49^. The main factor causing the age-related gamma enhancement might be maturation of inhibitory GABAergic neurotransmission that is protracted into early adulthood^26^. Changes in subunit composition of GABA receptors^50^, as well as increasing ‘on-demand’ GABA release^27^ produce a developmental shift toward stronger, more precise inhibitory currents, thereby resulting in highly coherent activity of principal cells at gamma frequencies^51^.

Developmental slowing of visual gamma oscillations found in the present study (Table 2, Fig. 4B) is generally in line with the results of Gaetz and colleagues^52^, who described changes in the frequency of visual gamma oscillations in a group of subjects aged 8–46 years. Inspection of the figure 2 of Gaetz et al^52^ suggests that the drop in gamma frequency in their participants occurred already during childhood. These authors, however, did not analyze the developmental changes separately in children and adults and attributed them to aging. Our study indicates that the most profound developmental decrease in gamma frequency occurs between childhood and adolescence and thus presumably reflects maturation of gamma-generating networks. This profound fall in gamma frequency might be causally related to a concurrent fall in the tonic excitability of inhibitory, most probably fast spiking parvalbumin positive (FS PV+), neurons^17,18,53^. There is convincing evidence for a developmental shift towards lower NMDA-mediated tonic excitation of the FS PV+ cells, as well as for a developmental reduction of the number of FS PV+ cells exhibiting excitatory NMDA-mediated currents^28^. On the other hand, there is an early developmental switch from depolarizing to hyperpolarizing actions of GABAA receptors^45–47^ followed by gradual increases in tonic inhibitory regulation of the FS PV+ interneurons during postnatal development^26^. These maturational transformations lead to subtler tonic excitability of inhibitory neurons and may well explain the strong (>10 Hz) and rapid decrease in the frequency of visual gamma oscillations that we observed between 7 and 15 years of age. Of note, the age-related decrease in expression of the NMDAR units (NR1, NR2a, NR2b) continues even in the mature brain - from young to middle-aged adulthood^44^ and may explain the more gradual decrease of visual gamma frequency in adults in our study.

When considering developmental changes in visual gamma oscillations, the constant magnitude of the velocity-related modulations of gamma frequency and power through childhood and adulthood is a remarkable finding. In light of the discussion above regarding the neural implications of gamma suppression, this constancy suggests that the global E-I balance in the visual cortex is efficiently regulated throughout a broad age range. Moreover, given the substantially higher gamma frequency in children as compared to adults (Tab. 1, Fig. 5B), the adjustment of the global E-I balance in the immature brain seems to be achieved at a higher level of excitability in both E and I neurons. The developmental constancy of gamma modulations is in line with an *in vitro* observation that the number of inhibitory synapses on the neuronal branches scales with the number of excitatory synapses during development, resulting in a relatively stable E/I ratio across several developmental stages^54^.

However, a few words of caution are warranted regarding the lack of a reliable developmental trend in gamma modulation parameters. According to animal studies, the maturation of neural processes underlying the E-I balance in auditory and visual cortices takes place during early sensitive periods of brain development^55^. All of our participants were older than 6 years of age, and it is possible that gamma modulation parameters could be age-dependent during infancy and toddlerhood, due to a continuous process of developmental fine-tuning of the E-I balance.

To summarize, we show that the bell-shaped changes in visual gamma power, that have been previously observed in the LFP of animals in response to increasing excitatory drive, are reliably reproduced in human MEG when the excitatory drive is modulated by the velocity/temporal frequency of drifting gratings. These non-linear power changes were paralleled by a nearly linear increase in gamma frequency. Based on previous animal and computational modeling studies, we speculate that the gamma suppression produced by sufficiently strong excitatory drive reflects the capacity of highly excited inhibitory neurons to control cortical gain in response to increasing input intensity. The suppression of gamma response at high velocities of visual motion appears to be an extremely robust effect, and its magnitude is scaled to the same range of values in neurologically typical children and adults, despite the drastic developmental changes in gamma response frequency and power. We speculate that gamma suppression parameters provide a quantitative and non-invasive measure of inhibitory-based gain control in disorders characterized by an altered balance between neural excitation and inhibition.

## Material and methods

### Data availability statement

The datasets generated and/or analyzed during the current study are available from the corresponding author on reasonable request.

### Participants

The inclusion criterion was the absence of neurological or psychiatric diagnoses reported by the participants (or their guardians). For the first experiment, 10 healthy volunteers (mean age 28.3 years, range 22–39 years, 5 males) were recruited among students and personnel at the Moscow University of Psychology and Education. The second experiment was performed in 50 children (all boys) and 27 adults (3 females). Children were recruited from schools in Moscow. Two children were later excluded from the MEG analysis because of technical reasons (excessive MEG artifacts or inability to follow instructions). Two more children (13 and 7 years) were excluded from the group analysis because post-hoc analysis revealed medical conditions (Arnold-Chiari malformation, minor motor ticks); their results are presented separately (Fig. 3). Twenty-seven adult subjects (3 females) were recruited in Gothenburg. One male (20 years-old) was later excluded from the MEG group analysis because of a post-hoc discovery of unreported epilepsy; his results are also presented separately (Fig. 3). The final sample comprised 46 children (6.7 - 15.5 years, mean 11.0, SD = 2.2) and 26 adults (19.2 -40.1 years, mean 27.9, SD=6.3).

The investigations were conducted in accordance with the Declaration of Helsinki and were approved by the Ethical Committee of the Moscow University of Psychology and Education or by the Gothenburg Regional Ethical Review Board (Regionalaetikprövningsnämnden i Göteborg). All participants provided their verbal assent to participate in the study and were informed about their right to withdraw from the study at any time during testing. Adult participants or guardians (for children) also gave written informed consent after the experimental procedures had been fully explained.

### Experimental task

The procedure used herein is similar to that used previously^25,43^. The stimuli were generated using Presentation software (Neurobehavioral Systems Inc., USA) and consisted of black and white sinusoidally modulated annular gratings with a spatial frequency of 1.66 cycles per degree of visual angle with an outer diameter of 18 degrees of visual angle (Fig. 1A). The gratings appeared in the center of the screen over a black background. In experiment 1, we presented 4 types of stimuli: a static grating and ones drifting to the central point at velocities of 1.2, 3.6, and 6.0°/s, (2, 6 and 10 Hz temporal frequency) referred to as ‘static’, ‘slow’, ‘medium’, and ‘fast’. In experiment 2, only the three types of moving gratings were presented. Each trial began with the presentation of a white fixation cross in the center of the display over a black background for 1 200 ms that was followed by the presentation of one of the four (in experiment 1) or the three (in experiment 2) types of stimuli chosen at random. After a randomly selected period of 1 200 - 3 000 ms, the movement stopped or the static stimulus disappeared. Participants were required to respond to the change in the stimulation flow (stop of the motion or disappearance of the static stimulus) with a button press. If no response occurred within the given response period (see *Supplementary information)* a discouraging message, “too late !”, appeared and remained on the screen for 2 000 ms, after which a new trial began. Stimuli were presented in three experimental blocks summing up to 90 presentations of each stimulus type. In order to minimize visual and mental fatigue, short (3–6 s) animated cartoon characters were presented between every 2–5 stimuli.

### MEG data recording

For adults in experiment 1 and children in experiment 2, neuro-magnetic brain activity was recorded at the Moscow MEG Centre (the Moscow State University of Psychology and Education) using a 306-channel detector array (Vectorview; Neuromag, Helsinki, Finland). For adults in experiment 2, MEG was recorded at the NatMEG Centre (The Swedish National Facility for Magnetoencephalography, Karolinska Institutet, Stockholm) using a similar 306-channel system (ElektaNeuromag TRIUX). The data was recorded with a band-pass filter of 0.03–330 Hz in Moscow and 0.1–330 Hz in Stockholm, digitized at 1000 Hz. The subjects’ head positions were continuously monitored during MEG recordings.

### MEG data preprocessing

The data was first de-noised using the Temporal Signal-Space Separation (tSSS) method^56^ and adjusted to the common head position using MaxFilter™ (v2.2) software. For further pre-processing, we used the MNE-python toolbox^57^ as well as custom Python and Matlab scripts. The de-noised data was filtered between 1 and 145 Hz. Independent component analysis (ICA) was used for correction of biological artifacts. The data was epoched from -1 to 1.2 sec relative to the stimulus onset. After rejection of artifacts (see *Supplementary information* for details) the average number of epochs in experiment 1 was 79.3, 79.4, 79.3 and 80.0 for the ‘static’, ‘slow’, ‘medium’ and ‘fast’ conditions, respectively. In experiment 2 there were on average 70.5, 70.8 and 70.5 epochs in children and 79.7, 79.3 and 78.0 in adults for the ‘slow’, ‘medium’ and ‘fast’ conditions, respectively.

### Time-frequency analysis of the MEG data at the sensor level

Planar gradiometers were used for the sensor-level analysis. We used discrete prolate spheroidal sequences (DPSS) multitaper analysis to estimate spectra in overlapping windows of 400 ms (shifted by 50 ms) at frequencies from 2.5 Hz to 120 Hz with a 2.5 Hz step and spectral smoothing of ±5 Hz around each center frequency. To decrease the contribution of phase-locked activity, the averaged evoked response was subtracted from each single data epoch.

The strength of sustained gamma response^29^ was estimated in dB in the 200–1200 ms post-stimulus period relative to the -900 to 0 ms period of pre-stimulus baseline. The individual gamma response frequency and power were then assessed at the posterior gradiometer’s pair with the maximal average response. Gamma response was considered ‘reliable’ if the post-stimulus increase in the peak power was significant at p<0.0001. (See *Supplementary information* for details).

To assess weighted gamma peak frequency we first found the absolute peak of gamma response for each subject and condition. As stated in the caption to Fig. 1, a frequency range of interest was established where the post-/pre-stimulus power ratio exceeded 2/3 of this peak value. The center of gravity of the power over this frequency range was used as the gamma peak frequency. The weighted gamma response power was calculated in dB, as 10*log_10_(post-stimulus/baseline), where post-stimulus/baseline is an average ratio over these frequencies.

### Time-frequency analysis of the MEG data at the source level

Source analysis has been performed for 23 adult participants in experiment 2 using FieldTrip software: http://www.ru.nl/neuroimaging/fieldtrip/ (see *Supplementary information* for details). For each subject, the brain MRI was morphed to the MNI template brain using linear normalization and a 0.6 mm grid. At the first step the time-frequency analysis was performed using DPSS multitapers centered at the sensor-defined subject/condition-specific frequency ±25 Hz. The DICS inverse-solution algorithm^39^ was used to derive common source spatial filters. In each participant/condition, the presence of a significant (p<0.05) brain cluster of post-stimulus increase in gamma power in the visual cortex was verified using bootstrap resampling source statistics (10000 Monte Carlo repetitions). At the second step, the ‘virtual sensor’ time series were extracted, and time-frequency analysis was performed. The gamma response parameters were then computed for the spectral average of 26 voxels adjacent to and including the one exhibiting the most significant post-stimulus increase in gamma power (using the same approach described above for the sensor space analysis). Details on the MEG analysis are given in the *Supplementary information*.

### Pupillometry

Changes in pupil size were regarded as indirect evidence of changes in excitatory drive as a function of visual motion velocity. We expected that an increase in excitatory drive with increasing velocity would be strong enough to affect pupil size. Pupil size was measured in 19 of the adult participants in experiment 2 using the EyeLink 1000 binocular tracker. We calculated the amplitude of the maximal pupil constriction (i.e., minimal pupil size) during the 0 to 1200 ms post-stimulus interval as a percentage of the change in average pupil size relative to the −100 to 0 ms pre-stimulus interval:

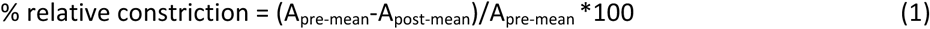

These values were computed separately for the slow, medium, and fast velocity conditions and then normalized to the average of the three conditions. Trials contaminated by blinks in the time window of interest (-100 to 1200 ms), as well as those preceded by cartoons and error trials were excluded from the analysis.

### Statistical analysis

We assessed the reliability of stimulus-related changes in gamma power in each subject and for each velocity condition using single epoch analysis. The pre- and post-stimulus power at the frequency of the maximal gamma response was compared using the Mann–Whitney U test.

Group analysis was performed using STATISTICA-13 software (Dell Inc, 2015). To test for the effect of the velocity condition on MEG parameters and pupil constriction magnitude, we used the repeated measures analysis of variance (rmANOVA). The Greenhouse-Geisser correction for violation of sphericity has been applied when appropriate. The original degrees of freedom and corrected p-values are reported together with epsilons (ε). Partial eta-squared (η^2^) was calculated to estimate effect sizes. ANOVA was also used to assess the effect of age group (i.e., adults and children). Spearman coefficients were used to calculate correlations between variables. For the correlation analysis presented in Fig. 6 we calculated normalized changes in response power for slow vs medium:

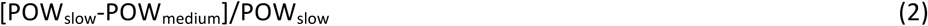
and medium vs fast:

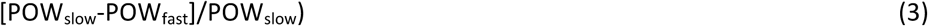

A more detailed description of the methods is provided in the *Supplementary information*.

## Acknowledgements

This work is financed by Torsten Soderberg Foundation (M240/13to CG) and the Russian Scientific Foundation (14–35–00060 to TS). Authors JFS and BR are supported by the Knut and Alice Wallenberg Foundation (grant 2014.0102), the Swedish Research Council (grant 621–2012–3673), and the Swedish Childhood Cancer Foundation (grant MT2014–0007). The author NH was supported by the LifeWatch Foundation. The author TS was supported by the Charity Foundation “Way Out”.

We heartily thank all participants and children families for their participation in this study. We are grateful to Daniel Lundqvist, Saideh Rajaei. Maria S. Davletshina, Anna V. Butorina and Anastasia Yu. Nikolaeva, for the technical support and/or help with data collection.

## Author contributions statement

E.V.O. and T.A.S. conceived the experiments. E.V.O., J.F.S., I.A.G., D.E.G., A.O.P., and B. R. conducted the experiments. E.V.O., T.A.S. and C.K. analyzed the results. E.V.O., T.A.S., J.F.S., N.H., C.G. and S.L. wrote the manuscript. All authors reviewed the manuscript.

